# Microglia complement signaling promotes neuronal elimination and normal brain functional connectivity

**DOI:** 10.1101/2021.03.31.437118

**Authors:** Senthilkumar Deivasigamani, Mariya Timotey Miteva, Silvia Natale, Daniel Gutierrez-Barragan, Bernadette Basilico, Silvia Di Angelantonio, Constantin Pape, Giulia Bolasco, Alberto Galbusera, Alessandro Gozzi, Davide Ragozzino, Cornelius T. Gross

**Author notes:** Current address: Institute of Science and Technology (IST) Austria, Klosterneuburg, Austria. **Classification:** Biological Sciences, Neuroscience.

## Abstract

Complement signaling is thought to serve as an opsonization signal to promote the phagocytosis of synapses by microglia. However, while its role in synaptic remodeling has been demonstrated in the retino-thalamic system, it remains unclear whether complement signaling mediates synaptic pruning in the brain more generally. Here we show that mice lacking the complement 3 receptor (*C3r*), the major microglia complement receptor, fail to show a deficit in either synaptic pruning or axon elimination in the developing mouse cortex. Instead, mice lacking C3r show a deficit in the perinatal elimination of neurons, both in the retina as well as in the cortex, a deficit that is associated with increased cortical thickness and enhanced functional connectivity in these regions in adulthood. These data demonstrate a preferential role for complement in promoting neuronal elimination in the developing brain and argue for a reconsideration of the role of complement in synaptic pruning.

## Introduction

Complement signaling is well documented to be a critical pathway for marking cells for phagocytic engulfment by macrophages in multiple organs across the mammalian body. The complement pathway is triggered by the binding of C1q (1, 2) to a variety of cell death markers presented on the cell surface of apoptotic cells and the subsequent enzymatic activation of a proteolytic cascade of C3, C4 and C5b-9 (3–5). The resulting complex of complement factors serves as a cell surface, or opsonization signal for recognition by macrophages expressing the complement receptor C3r (3). Other complement pathway components such as mannose binding lectin and properdin have also been shown to promote phagocytic uptake of apoptotic cells (6–8). The discovery that complement factors are expressed in the central nervous system and that microglia, the resident brain phagocytic cells, express C3r suggested that this pathway was also likely to contribute to neuronal phagocytosis and elimination (9, 10). The phagocytic elimination of neurons by microglia during brain development is well documented by histological studies in which apoptotic neurons were found surrounded by microglia (11–16) and by studies in which the pharmacological depletion of microglia resulted in an excess of neurons (12).

Nevertheless, evidence emerged showing that mice lacking *C3r* showed deficits in the developmental refinement of binocular retino-thalamic projections (17), a phenotype that suggested a role for complement in the phagocytic elimination of synapses or axonal branches, rather than whole neurons. This finding has important implications because the connectivity of the adult brain is the result of a more than two-fold overproduction of connections and their subsequent elimination during development, a process called synaptic or axonal pruning, depending on whether it involves the elimination of local synaptic specializations or the elimination of whole axonal branches, respectively (18, 19). For example, axonal pruning is well documented to occur in inter-hemispheric cortical projections with over 70% of connections being lost in the first six months after birth in primates (20) and similar levels of axonal pruning occuring in the postnatal rodent cortex (21, 22). Moreover, evidence points to a second wave of synaptic pruning during adolescence (23–25) that has been implicated in the etiology of schizophrenia (26–28). Thus, the identification of a molecular signaling pathway underlying these processes had major implications for the field and promised to open the door for the first time to their mechanistic study.

However, so far a role for complement signaling in synaptic or axonal pruning outside of the retino-thalamic system lacks strong evidence (29–31). Here we show that cortical structures in mice lacking C3r undergo apparently normal adolescent synaptic pruning and perinatal axonal pruning. However, axonal pruning of retino-thalamic projections was significantly reduced. Because axonal pruning in the retino-thalamic pathway is the result of retinal ganglion cell (RGC) death, rather than the selective elimination of their axonal projections, we hypothesized that complement signaling might exert an effect on neuronal connectivity via its role in promoting microglia-mediated cell elimination. Consistent with this hypothesis *C3r* mutant mice showed decreased neuronal elimination in the perinatal cortex and an associated increase in cortical cell number and synaptic connectivity in adulthood.

## Results

### Normal synaptic and axonal pruning in C3r knockout mice

Our initial experiments focused on determining whether mice with deficient complement signaling showed evidence of a failure in synaptic pruning as had been shown in the retino-thalamic system (17, 32). We chose to study mice lacking the complement 3 receptor (*C3r* knockout) because this mutant was previously shown to exhibit retino-thalamic pruning deficits and because it is the major complement receptor expressed by microglia, the primary phagocytic cell-type implicated in synaptic pruning in the mammalian brain (33–35). First, we monitored adolescent synaptic pruning by quantifying excitatory spine density in layer 5 neurons of the prelimbic cortex either before (postnatal day 30) or after (postnatal day 60) sexual maturity in the mouse (**Figure 1A**). As previously described (25, 36), a significant reduction in spine density was observed in all animals between P30 and P60, but no significant effect of genotype or interaction between time and genotype emerged (**Figure 1A-B**). These data suggest that adolescent synaptic pruning of principal cortical neurons proceeds normally in *C3r* knockout mice.

**Figure 1.**
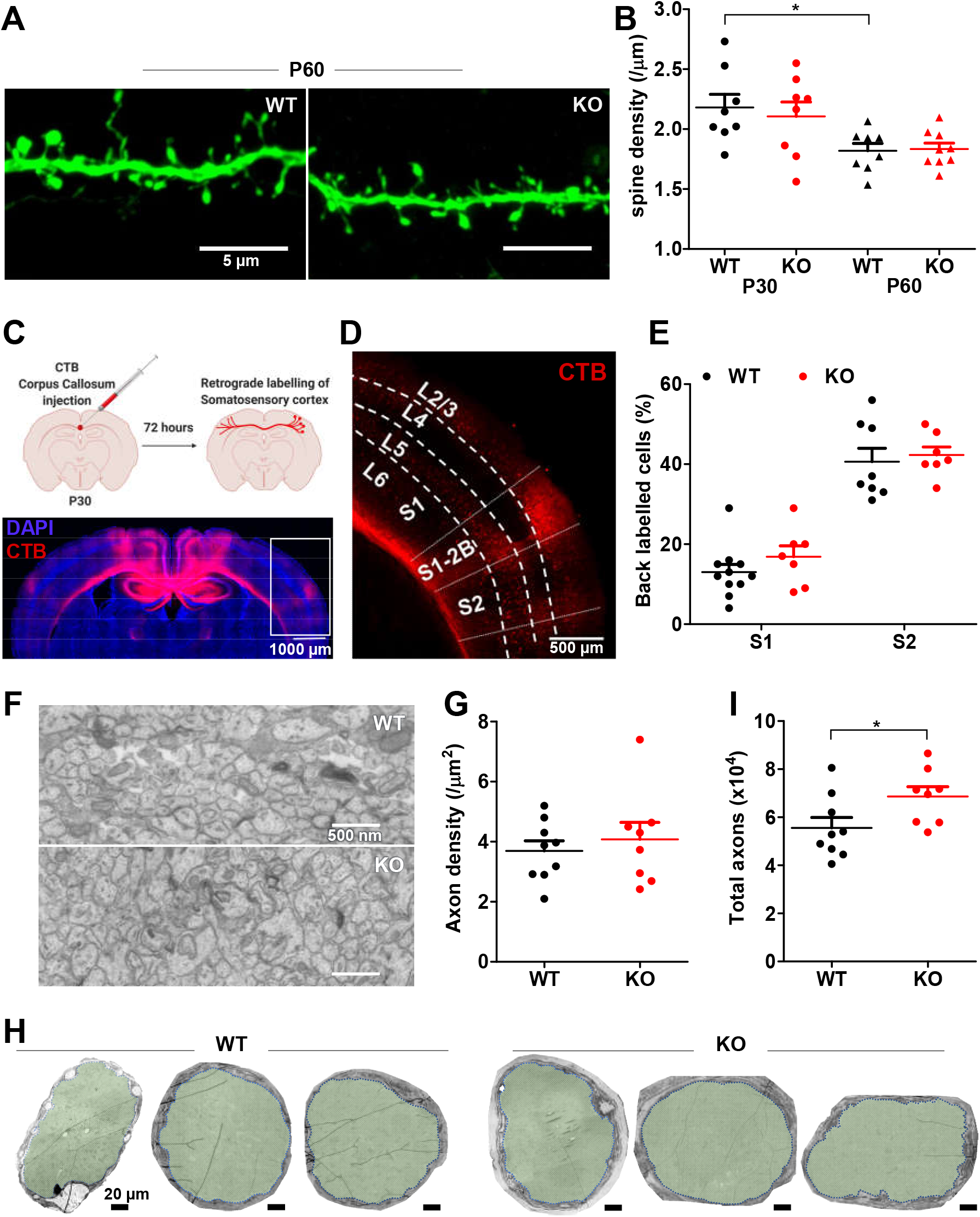
Normal synaptic and axonal pruning but deficient retinal ganglion cell axonal pruning in C3r knockout mice. (**A-B**) Pruning of dendritic spines during adolescence was examined in *C3r* knockout mice at P30 and P60. (**B**) Quantification revealed no synaptic pruning deficits in *C3r* knockout mice during adolescence (two-way ANOVA with Tukey’s posthoc test. main effect of time: F[1, 29] = 12.71, p = 0.001), main effect of genotype: F[1, 29] = 0.11, p = 0.737), time x genotype interaction: F[1, 29] = 0.25, p = 0.620). (**C**) Retrograde labelling was used to examine callosal axon pruning by direct injection of CTB into corpus callosum. (**D**) CTB Labelling of S1 and S2 somatosensory cortex regions showing callosally projecting neurons at postnatal day 30. (**E**) Quantification of callosally projecting neurons revealed no differences between wild-type and *C3r* knockout in the S1 and S2 regions of the somatosensory cortex (two-way ANOVA with Tukey’s posthoc test. main effect of region: F[1, 29] = 106.00, p < 0.001), main effect of genotype: F[1, 29] = 1.15, p = 0.292), region x genotype interaction: F[1, 29] = 0.18, p = 0.672). (**F-G**) Lack of *C3r* does not alter axon density in postnatal day 6 optic nerve (t-test, p = 0.554). (**H**) *C3r* knockout mice have a marginal increase in optic nerve diameter and (**I**) increase in total axon number (Unpaired t-test, p = 0.045). Each data point refers to an individual animal (mean ± SEM, * p < 0.05, ** p < 0.01).

Next, we quantified axonal elimination in inter-hemispheric cortical connections, a process that involves the pruning of more than 80% of callosal fibers in layer 4 of the primary somatosensory cortex in the first three postnatal weeks of mice (22). Local injection of the fluorescent retrograde tracer cholera toxin B (CTB) into fibers of the corpus callosum led to labeling of contralaterally-projecting cell bodies throughout the mouse cortex (**Figure 1C**). During the first postnatal week the vast majority of cell bodies in layer 4 of the S1 somatosensory cortex were labeled (22), while at postnatal day 30 less than 20% of soma showed labeling, demonstrating the efficient pruning of the majority of inter-hemispheric axons in this region (**Figure 1D-E**) (22). In the neighboring S2 somatosensory cortex pruning is less extensive and over 40% of soma showed persistent labeling (22) (**Figure 1D**). Quantification of the fraction of labeled cell bodies at postnatal day 30 revealed statistically indistinguishable pruning in *C3r* knockout mice and wild-type littermates in layer 4 of both S1 and S2 cortex (**Figure 1E**). These findings demonstrate that microglia-dependent complement signaling does not have a major role in mediating synaptic or axonal pruning in the developing mouse cortex.

### Deficient retinal ganglion cell axonal pruning in C3r knockout mice

Our initial findings suggested that synaptic or axonal pruning deficits in *C3r* knockout mice might be restricted to the retino-thalamic system. To test this possibility, we revisited the pruning phenotype in the visual system of these animals by directly quantifying the perinatal loss of retinal ganglion cell axons. Anatomical studies have shown that during the first postnatal week in rodents approximately half of retinal axons in the optic nerve are eliminated (37), while in carnivores and primates more than two-thirds are lost (20, 38, 39). Quantification of total axon numbers in electron micrographs of the prechiasmatic optic nerve at postnatal day 6 of *C3r* knockout and wild-type littermates (**Figure 1F**, **Figure S1A**) revealed a significant deficit in axonal pruning in the mutant mice seen as an increased total number of axons without a change in axon density (**Figure 1G-I**, **Figure S1B**). Assuming a 50% loss of axons in the mouse (37), these data suggested that microglia-dependent complement signaling is responsible for about one-quarter of axonal pruning in the retino-thalamic system. This phenotype is of a similar magnitude as the deficit in retino-thalamic projection enervation seen in these and other complement system knockout mice and suggest that microglia-dependent complement signaling has an essential role in axonal pruning in the early visual system.

### Microglia phagocytose apoptotic neurons in early postnatal cortex

However, retinal ganglion cell axonal loss in perinatal development is well documented to be accompanied by retinal ganglion cell death (40–43), opening the possibility that complement signaling in this system may mediate apoptosis, rather than synaptic or axonal pruning *per se*. Indeed, a recent study has documented a significant deficit in the loss of retinal ganglion cells in *C3r* knockout mice and the magnitude of the reported deficit is similar to the loss of axons we find (44). It is well documented that microglia seek out and engulf apoptotic neurons during brain development (12, 13, 45) and drugs that interfere with the phagocytic capacity of microglia result in a decrease in neural precursor death during cortical development (45). Two major phases of cortical apoptosis have been described. The first occurs in the subventricular proliferative zone across cortical development in close connection with cortical neurogenesis and is responsible for the death of a significant fraction of newly born cortical neurons (46, 47). Mutations in Caspase 9 that block cell death during this phase of cortical development are associated with macrocephaly, severe cortical malformations, and perinatal lethality (48). Given the viability and apparently grossly unaffected brain morphology of *C3r* knockout mice we thought it unlikely that complement proteins would be involved in the death of neuronal precursors. However, a second phase of cortical apoptosis has been described that occurs in the early postnatal period and involves the death of a subset of neurons that have successfully migrated into the cortical plate (11, 49–51). For reasons that at present remain unclear, this apoptotic phenomenon has been reported to be most frequent in medial cortical regions (49, 52).

To determine whether microglia might be implicated in the phagocytosis of apoptotic neurons during this second phase of cell death we carried out staining for apoptotic markers and microglia in the ACC of mice at postnatal day 5. A small, but significant number of cells dispersed across cortical layers showed immunostaining for the activated form of the apoptosis-associated factor Caspase 3 (aCasp3+) and in the majority of cases (26/33 cells, 78%) aCasp3+ cells showed condensed, pyknotic nuclei (**Figure 2A**). Nevertheless, only a minority of pyknotic nuclei expressed aCasp3+ (26/192 cells, 13.5%), suggesting that either caspase activation is not a prerequisite for pyknotic cell death, or that it represents a transient stage in the apoptotic process (**Figure 2D, Figure S2A-C**). A significant fraction of pyknotic cells were found to be engulfed by microglia as detected by immunostaining with the marker Iba-1 (**Figure 2B**) consistent with the hypothesis that microglia play a major role in the phagocytosis of this type of apoptotic postnatal cortical cell. Compared to aCasp3+ cells, pyknotic nuclei labeled with the live fluorescent sensor for extracellular phosphatidyl-serine (PS) modified lipids, PSVue (53–55), were preferentially found engulfed by microglia (**Figure 2C-E**). We interpret these findings to indicate that Caspase 3 is activated transiently in cells during early stages of apoptosis following which these cells progress to nuclear condensation, PS exposure, and identification by microglia for engulfment. Finally, we examined the identity of the engulfed pyknotic cells by repeating the co-labeling experiments in mice expressing the tomato fluorescent protein exclusively in cortical excitatory neurons (*Emx1*::Cre; *RC*::LSL-tomato; **Figure 2F**). About half of the engulfed cells could be confirmed as excitatory neurons (**Figure 2G**).

**Figure 2.**
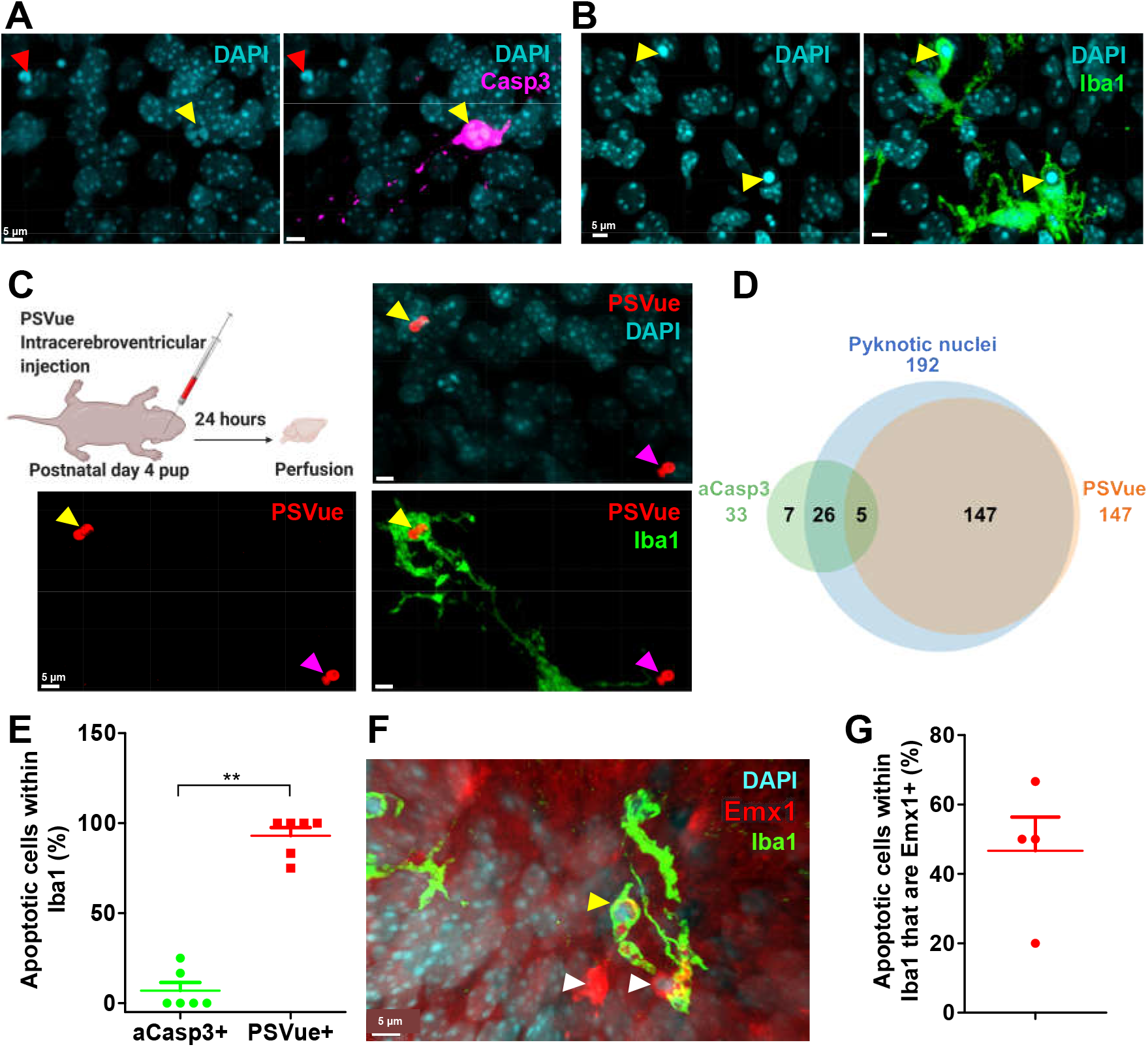
Microglia phagocytose apoptotic neurons in early postnatal cortex. (**A**) Pyknotic nuclei (red arrowhead) and apoptotic cells labelled with activated caspase-3 (yellow arrowhead) were found in early postnatal cortex. (**B**) Microglia phagocytose apoptotic cells (yellow arrowhead) in early postnatal cortex. (**C**) Apoptotic cells expose phosphatidyl-serine (PS) as labelled with PSVue (magenta arrowhead) and are phagocytosed by microglia (yellow arrowhead). (**D**) Quantification of pyknotic nuclei from anterior cingulate cortex (ACC) revealed that the majority of the nuclei are positive for PSVue but not activated caspase-3. (**E**) PS exposing cells are preferentially phagocytosed by microglia (Mann-Whitney test, p = 0.004). (**F**) The majority of pyknotic nuclei in ACC are of Emx1 lineage as seen by expression of tdTomato (white arrowhead) and they are phagocytosed by microglia (yellow arrowhead). (**G**) Cells of Emx1 lineage were found phagocytosed by microglia in extensively (One sample Wilcoxon signed-rank test, p = 0.097). Each data point refers to an individual animal (mean ± SEM, * p < 0.05, ** p < 0.01).

### Deficient neuronal elimination in C3r knockout mice

To determine whether complement signaling might play a role in the recognition and engulfment of apoptotic cells by microglia we quantified pyknotic cells and their engulfment by microglia in the anterior cingulate cortex (ACC) and somatosensory cortex of *C3r* knockout and littermate control mice at postnatal day 5. When compared to controls the density of pyknotic cells was significantly increased in *C3r* knockout animals in the ACC, but not in somatosensory cortex where apoptotic cell density was overall significantly lower (**Figure 3A**), consistent with the gradient of early postnatal cell death from medial to lateral cortical areas reported earlier (49). A smaller fraction of pyknotic cells were found engulfed by microglia in the ACC of *C3r* mutant mice when compared to controls, although this difference was not significant (**Figure 3B**). Overall, a significantly larger fraction of pyknotic cells were found engulfed by microglia in ACC when compared to somatosensory cortex, further strengthening the idea of a medial-to-lateral gradient (**Figure 3B**). We also observed a significant decrease in the number of phagocytic cups in the ACC of *C3r* mutant mice compared to control littermates, suggesting a reduced phagocytic capacity in the mutants and an overall lower phagocytic capacity in somatosensory compared to ACC (**Figure 3C**). Quantification of a Phagocytosis-Apoptosis Index (Iba-1+ pyknotic cells/ (total microglia*total pyknotic cells)) revealed a highly significant reduction in *C3r* mutant mice when compared to controls (**Figure 3D**). We also observed a small, but non-significant increase in the density of microglia in *C3r* mutant mice possibly reflecting an increase in microglia chemotaxis linked to reduced phagocytic capacity (**Figure S3**). Together these data suggest that microglia lacking C3r are less efficient at identifying or engulfing pyknotic cells and that this process is favored in medial versus lateral cortical structures.

**Figure 3.**
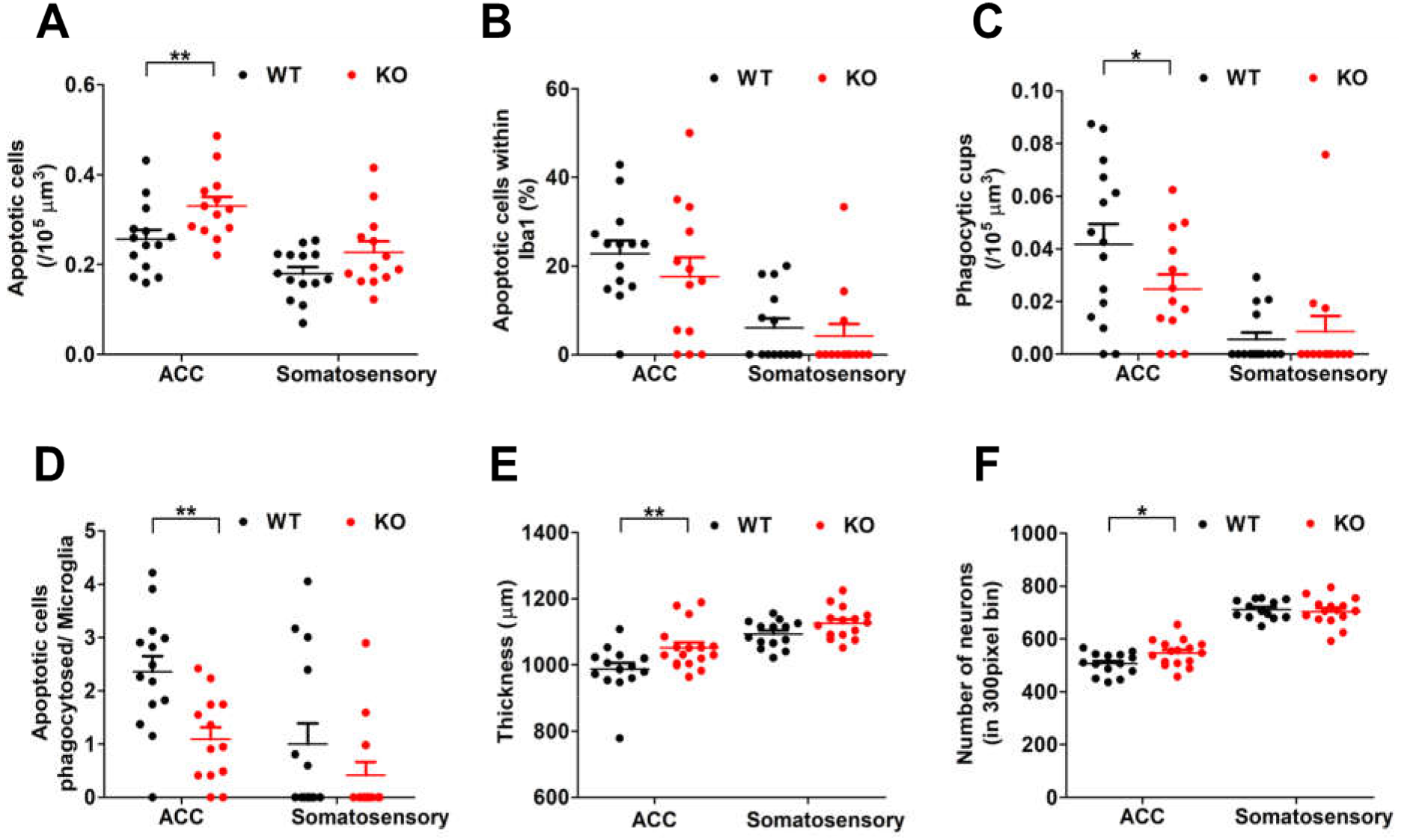
Deficient neuronal elimination in C3r knockout mice. (**A**) *C3r* knockout mice have increased density of pyknotic nuclei in the ACC but not in somatosensory cortex (two-way ANOVA with Tukey’s posthoc test. main effect of region: F[1, 50] = 23.19, p < 0.001), main effect of genotype: F[1, 50] = 10.66, p = 0.002), region x genotype interaction: F[1, 50] = 0.72, p = 0.399). (**B**) A smaller but non-significant fraction of pyknotic cells were found engulfed by microglia in the ACC of *C3r* knockout mice when compared to wildtype. However, a clear regional difference was observed (two-way ANOVA with Tukey’s post hoc test – main effect of region: F[1, 50] = 25.87, p < 0.001); main effect of genotype: F[1, 50] = 1.06, p = 0.307); region x genotype interaction: F[1, 50] = 0.19, p = 0.665). (**C**) *C3r* knockout mice have reduced number of phagocytic cups compared to wildtype littermates (two-way ANOVA with Tukey’s post hoc test – main effect of region: F[1, 50] = 49.74, p < 0.001); main effect of genotype: F[1, 50] = 4.24, p = 0.045); region x genotype interaction: F[1, 50] = 4.45, p = 0.040). (**D**) *C3r* knockout mice have reduced phagocytosis-apoptosis index in the ACC compared to control littermates (two-way ANOVA with Tukey’s post hoc test – main effect of region: F[1, 50] = 11.22, p = 0.001; main effect of genotype: F[1, 50] = 9.27, p = 0.003; region x genotype interaction: F[1, 50] = 1.26, p = 0.266). (**E**) Increased ACC cortical thickness is seen in adult *C3r* knockout mice (two-way ANOVA with Tukey’s post hoc test – main effect of region: F[1, 56] = 35.44, p < 0.001; main effect of genotype: F[1, 56] = 10.60, p = 0.002; region x genotype interaction: F[1, 56] = 1.96, p = 0.300). (**F**) Increased number of neurons in the ACC of *C3r* knockout mice (two-way ANOVA with Tukey’s post hoc test – main effect of region: F[1, 54] = 35.44, p < 0.001; main effect of genotype: F[1, 54] = 1.93, p = 0.170; region x genotype interaction: F[1, 54] = 227.09, p < 0.001). Each data point refers to an individual animal (mean ± SEM, * *p* < 0.05, ** *p* < 0.01).

The increased density of pyknotic cells in *C3r* knockout mice (**Figure 3A**) suggests that less efficient microglia phagocytosis might not necessarily be associated with a change in upstream processes that trigger cell death. Some studies have shown, however, that in the absence of phagocytic engulfment, pyknotic cells can be rescued from cell death (56, 57). To determine the outcome of an absence of microglia complement signaling on cell survival we measured cortical thickness and absolute cell numbers in the ACC and somatosensory cortex later in adulthood. Both cortical thickness and the absolute number of neurons was significantly increased in the ACC, but not somatosensory cortex in *C3r* knockout mice when compared to control littermates (**Figure 3E-F**). These findings argue for a role of microglia complement signaling as a limiting factor in postnatal cortical neuron elimination.

### Increased synaptic and functional connectivity in C3r knockout mice

Next, we examined whether the lack of microglia complement signaling during development might have long-term consequences on neural connectivity and synaptic function. First, we quantified spontaneous excitatory synaptic responses in principal neurons of *ex vivo* hippocampal slices (**Figure 4A**). A significant increase in spontaneous excitatory synaptic current (sEPSC) amplitude, but not frequency was observed in *C3r* knockout neurons compared to those from littermate controls (**Figure 4A, Figure S4A, C**). However, no significant difference in the amplitude or frequency of miniature excitatory synaptic currents (mEPSC) was detected between genotypes (**Figure 4B, Figure S4B, D**), although an increase in the relative difference in amplitude between spontaneous and miniature events (Δ amplitude = sEPSC-mEPSC) was found in *C3r* knockouts when compared to controls (**Figure S4E**), suggesting increased synaptic multiplicity. On the other hand, excitatory synaptic responses to local extracellular electrical stimulation were significantly reduced in *C3r* knockout neurons compared to those from control animals (**Figure 4C**). This apparent discrepancy suggests that while synaptic multiplicity is enhanced in a small subset of active neural connections, on average synaptic strength is reduced in the absence of complement signaling in microglia.

**Figure 4.**
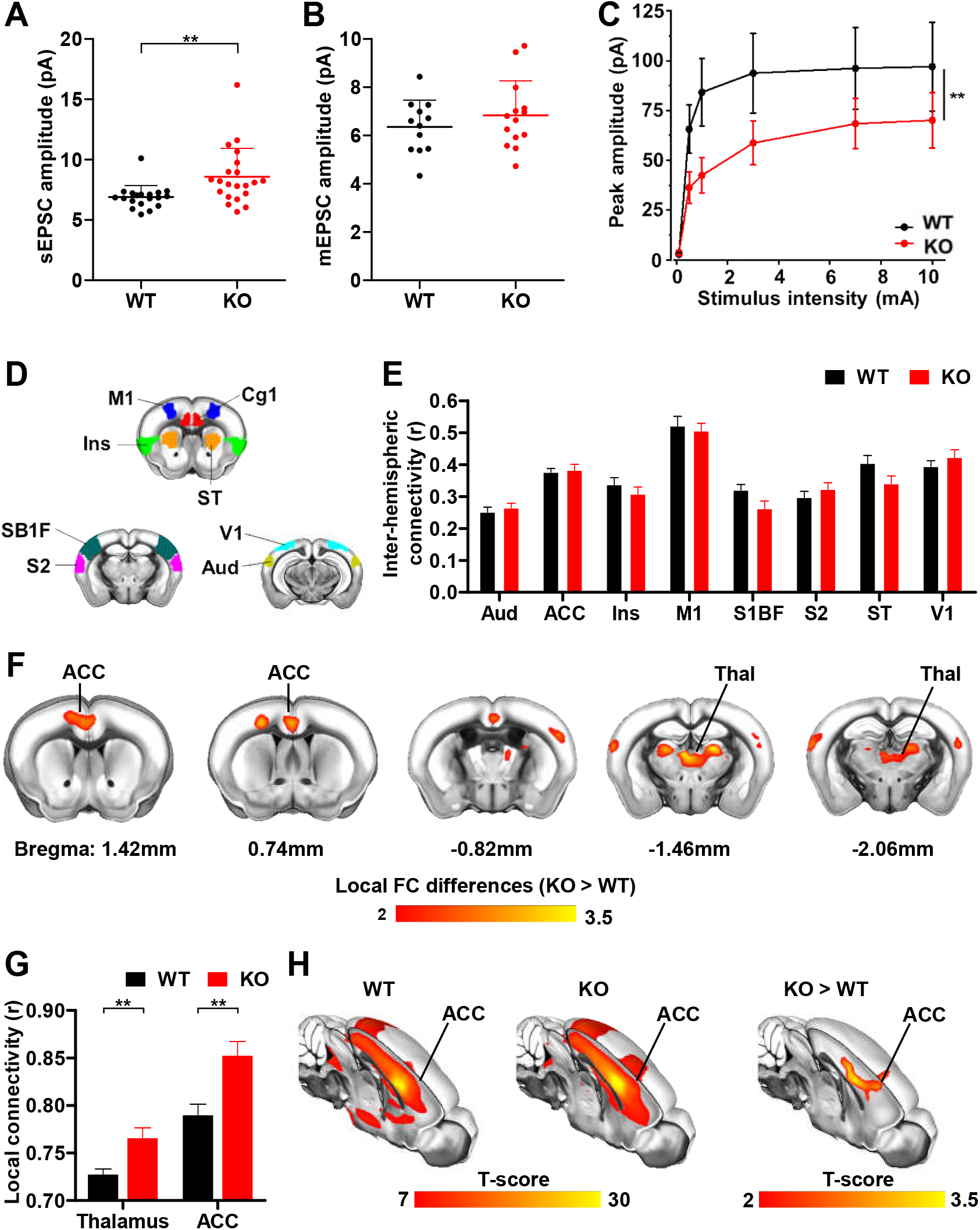
Increased synaptic and functional connectivity in C3r knockout mice. **(A)** *C3r* knockout mice at P40 showed significantly larger sEPSC than wild type littermates (unpaired t-test, p = 0.006). (**B**) On the other hand, no difference in amplitude (unpaired t-test, p = 0.66) of mEPSC were detected between the two genotypes. (**C**) *C3r* mutants exhibit reduced evoked responses than wildtype controls (two-way ANOVA with Bonferroni’s correction – main effect of genotype: F[1, 265] = 8.24, p= 0.004; main effect of stimulation intensity: F[5, 254] = 14.96, p< 0.001; stimulation intensity x genotype interaction: F[5, 265] = 0.44, p = 0.810) Each data point refers to an individual cell. (**D-E**) Group-level quantification of interhemispheric functional connectivity between pairs of mirror cortical regions, shows no alterations in long-range homotopic connections in *C3r* knockout mice. (**F**) Voxel-wise rsfMRI mapping of Local Connectivity alterations associated to *Cr3* mutation (unpaired t-test, p < 0.05, FWER corrected, cluster-defining threshold T=2). (**G**) Group-level quantification of Local connectivity in Thalamus and ACC (unpaired t-test, p < 0.01). Each data point refers to an individual cell. (**H**) Seed-based mapping of the default-mode network (DMN) spanned by voxel-wise connectivity to the ACC, and group-level differences showing the local effect of the mutation (mean ± SEM, * p < 0.05, ** p < 0.01).

Finally, we took an unbiased functional magnetic resonance imaging (fMRI) approach to quantify gross brain connectivity changes in the mutant animals. No significant differences in long-range functional connectivity were detected between *C3r* knockout and control littermates as measured by interhemispheric connectivity (**Figure 4D-E**). The absence of interhemispheric connectivity changes confirmed our anatomical work showing a lack of impact of microglia complement signaling on interhemispheric cortical axon pruning (**Figure 1C-E**). On the other hand, *C3r* knockout mice showed foci of increased local functional connectivity in midline cortical structures centered around the ACC as well as in dorsal thalamic regions (mediodorsal thalamus – MD), two brain regions known to be highly anatomically interconnected (**Figure 4F-G**). We next probed whether these focal local connectivity differences would be associated with long-range connectivity of the corresponding network systems by using a seed-based probing of the ACC (58). In keeping with our previous inter-hemispheric probing of fMRI connectivity, between group differences in fMRI connectivity were focal and short-ranged (**Figure 4H**). These findings confirm a long-term impact of microglia complement signaling that is specifically targeted to dorsal midline brain regions.

## Discussion

The discovery that complement signaling mediates the pruning of synaptic connections in the developing retino-thalamic system (32) opened the door to studies showing that microglia are important mediators of developmental synaptic refinement across the brain (17, 33, 59–61). However, it remained unclear whether complement had a role in synaptic pruning outside of the retino-thalamic system. This question gained increased importance with the discovery that common copy number variants in complement factor *C4* showed a dose-dependent association with risk for schizophrenia in human populations (31, 62, 63). Based on the demonstrated role of complement signaling in the retino-thalamic system in mice (17, 32) the hypothesis was put forward that complement signaling could have a role in neuronal pruning more generally. Several recent studies have found that overexpression of C4 in the developing mouse cortex impacts spine density and function (31, 64). However, loss of function mutations in the mouse *C4* gene did not show a phenotype in these studies (31), leaving open the question of whether complement mediates cortical pruning in the undisturbed animal. Here we confirmed that mice lacking the microglia complement receptor *C3r* do not show alterations in synaptic or axonal pruning, but instead show deficits in the elimination of apoptotic neurons, and that this has a long-term impact on brain functional connectivity in a manner that selectively affects Schizophrenia-associated brain regions.

### Microglia complement signaling in retino-thalamic pruning

Both the magnitude and direction of our finding of a significant increase in retinal ganglion cell axons in *C3r* knockout mice at postnatal day 6 (**Figure 1F-H**) are consistent with earlier data showing an increase in binocular retino-thalamic innervation at the same time point (17). Although we cannot rule out that *C3r* has an additional function in the refinement of retinal ganglion cell axonal or synaptic ramifications in thalamic target areas, we interpret our data as evidence that the thalamic phenotype of these mice is the secondary consequence of a deficit in the elimination of exuberant retino-thalamic axons. Moreover, as axon elimination in the retino-thalamic system is the result of retinal ganglion cell death (41, 43), we hypothesize that the primary physiological function of complement signaling in the developing eye is to promote microglia-mediated neuronal cell elimination. This hypothesis is supported by a recent study in which *C3r (44)* knockout mice were shown to have an excess of retina ganglion cells of a magnitude matching that of both our (**Figure 1F-H**) and other visual system phenotypes in complement mutants (17, 32, 62). It will be important to further test this hypothesis by examining whether blocking retinal ganglion cell apoptosis via independent signaling pathways (e.g. Caspase 9 (48, 65)) results in similar binocular refinement deficits.

### Microglia complement signaling and apoptosis

Programmed cell death during mammalian cortical development can be divided into two categories: a notable and wide-spread cell death that occurs in neural progenitors of the proliferative zones of the cortex during embryonic development, and a sparse cell death that occurs in mature neurons in the early postnatal cortical plate in a medial-to-lateral gradient (49, 52). While mutations that block cellular apoptosis non-specifically in the developing cortex result in a dramatic excess of neurons, macrocephaly, and perinatal lethality (48, 66, 67), the relatively modest excess of neurons we see in *C3r* knockout mice appear to be restricted to medial regions of the cortex (**Figure 3D-E**) suggesting that they may be the result of a selective deficit in the second category of cortical programmed cell death. This hypothesis is supported by the reduced number of pyknotic cells found engulfed by microglia in layer 2/3 & 5 of the ACC, but not somatosensory cortex at postnatal day 5 (**Figure 3B**) and a concomitant overall increase in pyknotic cells at this time point (**Figure 3A**). It should be pointed out, however, that the engulfment of pyknotic cells by microglia appears to be only partially impaired in *C3r* knockout mice, a finding that suggests that complement signaling is one of multiple pathways promoting microglia phagocytosis and is consistent with the relatively minor phenotypes reported in the brain of these mice (17, 62). Complement deposition preceding microglial phagocytosis has been observed in both apoptotic and newborn cells (9, 68). Our data are potentially consistent with those of at least one other study that reported a deficit in aCasp3+ cells in the early postnatal hippocampus of *C3r* knockout mice at this stage (13). Interestingly, intraventricular treatment with a C3r inactivating antibody one day prior to aCasp3+ immunostaining phenocopied the knockout, demonstrating that the deficit in apoptosis was the result of an acute, postnatal impairment of microglia function (13). Unlike in this study, however, we observed only a mild, non-significant decrease in aCasp3+ cell number in *C3r* knockout mice (**Figure S3B**) and an increase in pyknotic cells and a decrease in their engulfment by microglia (**Figure 3A**). This phenotypic discrepancy could be the result of the different tissues examined or could it be a compensatory response to decreased phagocytosis of apoptotic cells in the mutants.

### Medial-to-lateral phenotypic gradient

Our observation of anatomical and functional imaging phenotypes in *C3r* knockout mice that were restricted to the anterior dorsal medial cortex (**Figure 3A** and **3D-E, Figure 4H-I**) are consistent with the reported gradient of early postnatal neuronal apoptosis along the medial-to-lateral axis (49, 52), a finding that was confirmed in our data (**Figure 3A**). We noted that despite similar densities of microglia in ACC and somatosensory cortex (**Figure 3C**), a significantly smaller fraction of pyknotic cells were found engulfed by microglia in the more lateral region (**Figure 3B**) suggesting that the gradient may be the consequence of a difference in phagocytic capacity of microglia. Transcriptional profiling of microglia along this gradient might reveal clues about the molecular signaling that regulates their early postnatal cell engulfment program. It is striking that the regions of cortex most affected in the complement mutant are those shown by unbiased clustering analysis of rodent anatomical tract tracing studies to be hubs that support executive function, planning, and self-awareness (69). It remains to be determined whether this gradient is relevant for the link between copy number variation in complement factor *C4* and risk for Schizophrenia (62).

### Long-term impact of microglia complement signaling

Although we cannot at present be certain that the changes in cortical thickness (**Figure 3D-E**), synaptic function (**Figure 4A-C**), and functional imaging (**Figure 4H-I**) observed in adult *C3r* knockout mice depend entirely on deficits in early postnatal cell elimination, the matching medial-to-lateral gradients of postnatal microglia phagocytosis (**Figure 3B**), cortical thickness, and functional imaging phenotypes suggests that they may be causally related. Critically, the data argue that the increase in cortical thickness and neuronal number in the adult ACC is a direct consequence of a failure of microglia lacking C3r to efficiently engulf pyknotic cells in this structure during the early postnatal period. Such a causal link, however, requires that in the absence of microglia engulfment pyknotic neurons can reverse their nuclear condensation phenotype and survive to adulthood. Such a reversal of commitment to apoptosis has been described in cultured cells and can include the transient expression of activated caspases, for example (70–72). Alternatively, the increase in cortical neurons in *C3r* knockout mice could be driven by a signal emitted by mutant microglia that reduces the number of neurons that commit to apoptosis. Resolving these mechanisms will require precise measurement of the dynamics of apoptosis and microglia phagocytosis in the developing cortex.

Our results argue for an important effect of the early programmed elimination of neurons on synaptic connectivity in adulthood. To the best of our knowledge such a link has not been explored previously and it is not immediately clear how the number of neurons could affect synaptic connectivity, other than more neurons providing more synaptic sources and targets. Our electrophysiology data show that the amplitude of spontaneous EPSC is increased in *C3r* knockout mice, but evoked responses are diminished. We interpret this to indicate that synaptic multiplicity – the number of synaptic sites made by one neuron on another – is increased in a minority of synapses that are spontaneously active in the *ex vivo* slice preparation, but that in the majority of remaining synapses strength is reduced. More work will be necessary to understand the intermediate steps in synaptic maturation across postnatal development that are affected in C3r knockout mice, leaving open the alternative possibility that C3r signaling might also have an impact on synaptic maturation that is independent from its effect on neuronal phagocytosis.

In summary, we have presented evidence for a role of complement signaling in promoting the developmental elimination of neurons by microglia selectively in the ACC and shown that this process is required to achieve normal functional connectivity in adulthood. We failed to find evidence for a role of microglia complement signaling in synaptic or axonal pruning, opening the possibility that complement mutant mice demonstrate deficits in synaptic refinement as a secondary result of a deficit in the elimination of neurons rather than in axonal or synaptic pruning. These findings call for a reevaluation of the role of complement in brain development and demand further work to understand how neuronal apoptosis and phagocytosis can be regulated in a region-specific manner to shape adult brain connectivity and function.

## Materials & Methods

Animals, immunostaining, electron microscopy, image quantification, surgical procedures, analysis of spine morphology, in vitro electrophysiology, resting state fMRI and statistical analyses are described in SI Appendinx. Experimental procedures were approved by the Institutional Animal Care and Use Committees of European Molecular Biology Laboratory and Italian Ministry of Health.

## Acknowledgments

We thank EMBL Heidelberg Electron Microscopy facility for processing and imaging of optic nerve, Robert Prevedel for co-mentorship, Sarah Kaspar for help with statistical analysis and Francesca Zonfrillo, Claudia Valeri, Roberto Voci, and Valerio Rossi for mouse husbandry. We thank the EMBL Rome Microscopy Facility, Animal Facility, Histology Facility, Genetic and Viral Engineering Facility, Gene Editing and Embryology Facility, for their support throughout this project.

## Funding

The work was supported by EMBL, the European Commission under the H2020-MSCA EI3POD COFUND fellowship program (Grant agreement ID: 664726), and a NARSAD Young Investigator Award from the Brain and Behavior Research Foundation (Grant No:28355) to S.D. A.Go acknowledges support from the European Research Council (ERC)under the European Union’s Horizon 2020 research and innovation programme (#DISCONN; no. 802371 to A.Go.), the Brain and Behavior Research Foundation (NARSAD; Independent Investigator Grant; no. 25861), the Simons Foundation (SFARI 400101), the NIH (1R21MH116473‐ 01A1) and the Telethon foundation (GGP19177).

## Author contributions

Experiments and data analysis were carried out by S.D, M.T.M, S.N, G.B. Codes for Stardist training and analysis were written by C.P. B.B, S.D.A and D.R performed and analyzed the slice physiology experiments. D.G.B, A.G, A.Go carried out the fMRI experiments. S.D. and C.T.G. designed the experiments, C.T.G supervised the project and together with S.D. conceived the project and wrote the manuscript.

## Competing interests

Authors declare that they do not have any conflict of interest.

## Data availability

Data were not deposited to any database repositories. All data needed to evaluate the conclusions in the paper are present in the paper and/or the Supplementary Materials. The data supporting the findings are available within the manuscript. Codes used for analysis of neuronal quantitation have been deposited publicly in Github.

## Notes

### Competing Interest Statement

The authors have declared no competing interest.

