## Supplemental Data for "Microglia complement signaling promotes neuronal elimination and normal brain functional connectivity"

#### SI Materials & Methods

##### *Animals*

All mice tested were obtained by internal colonies from the European Molecular biology laboratory. Mice were maintained in temperature and humidity-controlled condition with food and water provided *ad libitum* and on 12-h light-dark cycle (light on at 7:00). *C57BL/6J* mice were obtained from local EMBL Rome colonies. The following transgenic mice lines were used: *Thy1*::EGFP-M (1) (Jackson Laboratory stock 007788) and *Rosa26-CAG::loxP-STOP-loxP-tdTomatoWPRE (2)* (Jackson Laboratory stock 007905), CD11b-deficient mice (3) (*C3r* or *Itgam*) (Jackson Laboratory stock 003991), *Emx1::Cre (4)*. Thy1::EGFP animals were bred with *C3r* mice to generate double transgenic mice. Animals homozygous for *Thy1::*EGFP and heterozygous for *C3r* were bred to get animals of the desired genotype. Heterozygote *C3r* animals were bred to obtain WT controls and KO animals. For all the experiments littermate WT and KO were used wherever possible. Both males and females were used indiscriminately. All experiments were performed in accordance 91 with EU Directive 2010/63/EU and under approval of the EMBL Animal Use Committee 392 and Italian Ministry of Health License 541/2015-PR to C.G. The fMRI experiments were conducted in accordance with EU 86/609/EEC, DL 116, January 1992 and the Guide for the Care and use of Laboratory Animals of the National Institutes of Health. All surgical procedures were performed under anesthesia.

##### *In vivo PSVue labelling*

The activated PSVue 550 was prepared according to manufacturer's instructions (Molecular Targeting Technologies) using Zinc Nitrate and diluted to 1mM using sterile water. For *in vivo* labelling of apoptotic cells, postnatal day 4 pups were anesthetized by placing on crushed ice for 2-3 mins. Animals were then placed on a custom stage and the head was illuminated with a fibre optic light source. 1µl of activated PSVue was injected into the lateral ventricles ($2/5^{th}$ of the distance from the lambda to eye) using a 32G Hamilton syringe and the syringe was left in place 20-30 secs after injection. The pups were placed under a warm lamp and returned to the dam after recovery. 24 hours after injection the pups were perfused intracardially with PBS and 4% PFA in the Phosphate buffer. The brains were post-fixed in 4% PFA overnight and transferred to 30% Sucrose for cryoprotection

##### *Immunostaining*

For immunostaining of cryostat sections (40µm or 30; Leica Microsystems) were incubated with blocking buffer (1x PBS + 2% BSA + 0.3% Triton X-100) for 2 hours followed by overnight incubation at 4°C with primary antibodies diluted in blocking buffer. The following antibodies were used: Goat anti-Iba1, Wako 011-27991, 1:250, Rabbit anti-activated caspase-3, Cell Signaling Technologies, 1:500, Rabbit anti-Iba1, Wako 019-19741, 1:500, Mouse anti-NeuN, Millipore MAB377, 1:200. For spine density analysis, Thy1::EGFP labeled Vibratome sections (50µm; Leica Microsystems) were blocked and then incubated overnight at 4°C with primary antibody (Chicken anti-GFP, Aves Lab 1:500) diluted in blocking buffer. Following incubation with primary antibody the tissues were washed in PBS, blocked and incubated with fluorophore conjugated secondary antibodies (Life Technologies) for 2 hours at room temperature. The tissues were mounted with MOWIOL or Prolong (Life Technologies).

##### *Electron microscopy and quantification of optic nerve*

Mice at postnatal day six, from five different litters, were deeply anesthetized with Avertin (400 mg/kg, i.p.; Sigma-Aldrich) and slowly (less than 1ml/min) perfused transcardially with 1x PBS and then 4% paraformaldehyde plus 2.5% Glutaraldehyde in 0.1 M phosphate buffer (PB), pH 7.4. Eyes were promptly extracted and the part of optic nerves attached (preceding the optic chiasm) dissected and placed in post-fixation buffer overnight (4% paraformaldehyde plus 2.5% Glutaraldehyde in 0.1 M phosphate buffer, pH 7.4). Any nerves that showed signs of damage were discarded. Samples were then shipped to Electron Microscopy Core facility site in EMBL-Heidelberg, for imaging, in 0.1M PB solution containing 0.5x fixative and with no air to avoid oxidation. Samples were then prepared for imaging: with 1% $OsO_4$/1.5% Potassium Ferrocyanide for 1h on ice, then with 1% $OsO_4$ in Sodium Cacodylate buffer 0.1M pH 7.4 on ice followed by 10x rinse in water and then stained in with 1% aqueous solution of Uranyl Acetate overnight at 4°C. Samples were then dehydrated with increasing concentration of Ethanol and after 100% Ethanol they were infiltrated in 3:1, 1:1 and 3:1 acetone:resin for 2h each step, and left in 100% resin overnight rotating. Samples were embedded into molds the next morning. Durcupan embedding was carried out in a flat orientation within a sandwich of ACLAR® 33C Films (Electron Microscopy Science) for 72h at 60°C. The samples were then cut and remounted on EPON blocks for sectioning.

Sections 90 nm thick were collected in the middle region of the optic nerve extracted (by avoiding the damaged extremities) and post-stained for 3 min with lead citrate. Images were acquired on a JEOL JEM-2100 Plus electron microscope at 120 kV with the SHOTMEISTER (JEOL software) and then collected and shared with LLP Viewer software. Montages were taken at 1200x and 5000x magnification.

The regions of interest selected along the vertical midline, over the longer axes of optic nerves, were exported in Fiji with a fixed width of 5.8 μm and axons were manually counted. Axons were identified as rounded white/light-grey objects surrounded by a darker border and no distinction was made among myelinated (very low percentage) and unmyelinated axons. The number of axons was then divided for the size of area exported (axons/unit area) and to obtain the total number of axons per optic nerve the axons/unit area was normalized for the whole area. The whole area of the optic nerves was manually drawn and measured in Fiji by carefully excluding surrounding immune cells and blood vessels.

##### *Image Analysis*

###### Spine density analysis

For spine density analysis, immunostained *C3r KO; Thy1::*EGFP and control brain sections corresponding to prelimbic and cingulate cortices (Bregma: 1.78  to 1.34) were imaged on a TCS SP5 resonant scanner confocal microscope (Leica Microsystems) with a 63x/1.4 oil immersion objective at 48 nm lateral resolution and  a z-step of 130 nm. Spine density quantitation was performed as described earlier (5). Briefly, images were deconvolved using Huygens Professional software (Number of iterations<50, quality change threshold – 0.1, theoretical point spread function.  Image J software (NIH) was used for semi-automatic quantification of spine density.  8-bit maximum intensity projections were used and only lateral spines were analyzed. Signal intensity was measured in the dendritic shaft and used for normalization across all datasets. Background signal was measured outside the dendritic shaft and 1.4 times the background signal was removed. Images were automatically thresholded using the Huang algorithm. The image was inverted and watershed algorithm was applied followed by clearing of the dendritic shaft and neck. Spine number subsequently analyzed using particle measurement, after sphericity (>0.3/1) and size (>0.005μm$^2$) thresholding to avoid false positives. Spine density was calculated by normalizing the number of spines to the length of the dendritic branch imaged. On average 7 dendritic branches and 780μm of dendrite was analyzed per animal. Animals from ten different litters were used for spine density analysis.

**Quantification of pyknotic cells and microglial phagocytosis**

Quantification of pyknotic cells and microglial phagocytosis was analyzed on cortical tissues from postnatal day 5 mice. Stained presumptive cingulate and somatosensory cortical regions were imaged on an Ultraview Vox Spinning Disk Confocal (Perkin Elmer) equipped with an EMCCD Camera (Hamamatsu C9100-50). Initially, the entire slide was scanned at 20x and the region of interest was imaged at higher magnification 60x/1.4NA oil immersion objective with a z-resolution of 0.3µm. Similar imaging parameters were used for both wild-type and knock-out animals. All the analysis was carried out blind to the genotype and phenotype. Quantification of pyknotic cells was performed using Imaris 8.0 or Imaris Viewer (Oxford Instruments) by converting the files to native Imaris file format. Pyknotic nuclei were identified by their distinct bright, condensed nuclei. All the engulfment events were manually confirmed by going through the Z-stack. For quantification of microglial cells, only Iba1 positive cells with their soma clearly visible in the field were included. On average 8 fields were analyzed per animal for each brain region. Pyknotic nuclei, phagocytic cups and microglial count were normalized to the volume of tissue used for quantification. Similar volumes were used for quantification of both wildtype and knock-out: ACC total volume: $99.2x10^6$ $um^3$ vs $89.7x10^6$ $um^3$ (WT vs KO) and somatosensory cortex total volume: $86.9x10^6$ $um^3$ vs $80.6x10^6$ $um^3$ (WT vs KO). Phagocytosis-Apoptosis index was calculated as follows = (proportion of pyknotic nuclei inside microglia)/(density of pyknotic nuclei * density of microglia). Animals from eight different litters were used for this analysis.

**Quantification of Caspase3 cell density**

For quantitation of caspase-3 density in *C3r* knockout and control mice, immunostained postnatal day 5 brain sections were imaged on an Ultraview Vox Spinning Disk Confocal (Perkin Elmer) equipped with an EMCCD Camera (Hamamatsu C9100-50). Initially, the entire slide was scanned at 4x and regions of interest were imaged using 40x/1.3NA oil immersion objective with a z-resolution of 0.5µm. Number of aCasp3+ cells were normalized to the volume of the tissue imaged.

**Cortical thickness analysis**

For cell counting and cortical thickness measurements coronal sections of 30 µm from postnatal day 120 mice were used. Animals from fourteen different litters were used for cortical thickness and cell count measurements. The sections were stained for NeuN and DAPI and imaged on an inverted fluorescence microscope (THUNDER Imaging System, Leica Microsystems) using 20x/0.8 NA objective. The analysis was performed using Fiji/ImageJ, each image was rotated properly to be straight. NeuN channel was used to visualize the white matter structure which was used to identify the reference point for the measurements. Coronal sections from 1.53 mm to 1.21 mm were used for ACC analysis. For measurement of ACC thickness, first, a 180° line was drawn and then moved down until it touches the forceps minor of corpus callosum (fmi). The point in which the line touches the fmi for the first-time structure was used as a reference point. The ACC thickness was measured by drawing a 180-degree line from the reference point until the end of layer 1. Data reported is an average of 3-4 different coronal sections for each animal.

For somatosensory cortex analysis, coronal sections from -1.46 mm to -1.57 mm were used. The parameters to measure cortical thickness in somatosensory cortex was experimentally determined to perform the measures always in the S1 region. For measuring the thickness of Somatosensory cortex, as a first step, an $180^0$ line was drawn and then rotated at 45 degrees, this line was moved up until it touched for the first time the hippocampus. The point in which the 45-degree line intercepts the beginning of the cortex was used as a reference point from which the 60-degree line was drawn, if the measurement is performed on the right side, and 120 degrees if on the left side. This line was extended until the end of the layer 1 and the thickness of the cortex was measured. Data reported is an average of 3-4 different coronal sections for each animal.

***Automated cell count using StarDist***

The images used for automated cell counting were obtained on an inverted fluorescence microscope (THUNDER Imaging System, Leica Microsystems) using 20x/0.8NA objective using Z-step of 0.85µm and a lateral resolution of 0.41µm. Image acquisition was controlled using LAS X software. All the sections were imaged for NeuN and DAPI channels and similar imaging conditions were used for all the animals. The images were computationally cleared using Large Volume Computational Clearing algorithm (LVCC), a proprietary package of LAS X (Leica Microsystems) before proceeding with counting. LVCC processing helps in reducing the out of focus blurs and background signal.

StarDist is a deep supervised machine learning tool developed for automated cell prediction (6). The method is suited for objects with a star-convex shape. When training the StarDist model, the images in the training set are randomly split into training data (90%) and validation data (10%). The validation data is used during the StarDist training to monitor the progress of the training and to determine the best combination of parameters of probability threshold and overlap threshold that allows the best prediction in the validation data. StarDist was trained using the scripts available at https://git.embl.de/grp-bio-it/ai4ia/-/tree/master/stardist. All the models were exported from StarDist using Tensorflow version 1.14 and Tensorflow version 1.14.0 was used in Fiji/ImageJ as well. For each model developed the following parameters were determined

- *Probability/Score Threshold* – Higher threshold results in fewer segmented objects, thus reducing chances of false positives.
- *Overlap Threshold* – Higher threshold allows segmented objects to overlap substantially. Allows for segmentation of cells in a crowded field.

**Image annotations with QuPath and StarDist training**

QuPath v0.2.0-m11 software was chosen for the cell annotations used for all StarDist models. Detailed cell annotations were performed with brush modality only in NeuN channel but DAPI channel of the same image was used for confirmation. The StarDist approach learns how to predict cells using geometric parameters as the distance from the center of the cell to the border of the cell. For the StarDist algorithm, every single signal needs to be annotated even if it is from a cell that is partially visible because for example on the border of the image. Taking this into consideration, annotation was performed in a 512x512 pixels square positioned inside the 600x600 pixels image. Using this approach, it was possible to fully annotate borderline cells even if part of the cell body is going outside the 512x512 pixels square, this approach allows us to maintain the real geometric distance from the center to the border of the cell.

A pipeline outlining development of Stardist based image quantitation is described in **Figure S5A**. We started attempting using a pre-trained model 'Versatile (fluorescent nuclei)'. However, we observed high levels of relative error (%). Then we switched to custom user-trained models. The training was performed using 10 NeuN channel images with 512x512 pixel size from two brain regions: ACC and somatosensory cortex. After training with images from ACC (called 'C cortex' model) and somatosensory cortex (called 'SS cortex' model) we found that the model trained with images from somatosensory cortex ('SS cortex') performed best in automated counting in both ACC and somatosensory cortex regions. In some of the training Data Science Bowl (DSB) was included. DSB is a dataset with 37,333 manually annotated nuclei in 841 2D images from more than 30 experiments across different samples, cell lines, microscopy instruments, imaging conditions, operators, research facilities, and staining protocols. The annotations were manually made by a team of expert biologists (7). Inclusion of DSB pre-training data did not improve the performance of either 'C cortex' or 'SS cortex' model. Training the 'SS cortex' model with DAPI data did not improve the performance of the model. In fact, it resulted in poor performance of the 'SS cortex' model (relative error (%) 4.01±0.30 vs 41.31±20.9 without vs with DAPI) during validation of images from Somatosensory cortex. To improve the performance of the 'SS cortex' model and to eliminate any randomness that might contribute to the performance of the model, using the same training set used in the 'SS cortex' model, 10 different re-trainings were performed. In each training, 10 % of the images were randomly selected and included in the internal validation. Of these new ten re-trained models 'SS cortex re-trained 5' show a decrease of relative error (%) from 4.01±0.30 to 3.18±0.37 ('SS cortex re-trained 5' vs 'SS cortex').

*Absolute error* = |Number of cells predicted by StarDist − Number of cells obtained by average of 3 counts by manual method|

*Relative error in %* = (Absolute error / Number of cells obtained by average of 3 counts by manual method) *100

**Manual validation of all StarDist models**

For manual validation, images of 600x600 pixel from the NeuN channel five each from the ACC cortex and somatosensory cortex were used. For each image, manual counting was performed 3 times and an average of the three values was used as the final measure of the cell count and calculation of relative error (%). The StarDist model was applied to each image and then the result of the predictions was quantified. For each prediction, the relative error (%) was calculated.

#### Application of the best StarDist model

The best model 'SS cortex re-trained 5' was applied to the images using StarDist2D plugin in Fiji/ImageJ, entering manually the parameters of probability threshold and overlap threshold. The model was applied to a 2D z-section chosen with the following criteria: 1) section with the highest intensity; 2) section in which cells are well visible and in focus. In the chosen z-section the analysis was performed in a rectangle drawn with the following criteria: 1) the first dimension was fixed at 300 pixels; 2) the second dimension corresponds to measure of cortical thickness. The Probability/Score Threshold used was 0.45561 and the overlap threshold was set to 0.40000. All the predictions by StarDist were manually verified to remove any false positives or false negatives. The data presented is an average of two sections analyzed per animal.

#### *Retrograde labelling of Callosal projection neurons*

#### Stereotaxic surgery and retrograde labelling

CTB back-labelling and quantitation was performed as described by De Leon Reyes et al., 2019 (8) with some modifications. Mice of age Postnatal day 30 were anesthetized with 4% Isoflurane and placed in a stereotactic frame (RWD Life Sciences); isoflurane in oxygen (1-2%) was administered to maintain anesthesia. The skin was incised and the skull surface was exposed. The skull was trepanated using a dental drill. Using a glass capillary (tip diameter – 30um) 500nl of CTB 647 (0.5% in Phosphate-buffered Saline, Life Technologies) was pressure injected into the Corpus Callosum (Co-ordinates: AP =-1.4, ML =0.70, and DV =-1.70 with an angle of $18^0$) at the rate of 50nl/per minute. The capillary was left in position for 10-15 minutes after injection and then retracted. Animals received a Carprofen (Rymadil 5 mg/kg, subcutaneous injection) as surgical analgesia. After allowing CTB migration for 72 h post-surgery mice were transcardially perfused with 4% PFA in 0.1M Phosphate buffer.

#### Imaging and analysis of CTB labelled cells

Brains were removed from the skull and left to postfix overnight in 4% PFA at 4°C. Brains were cryoprotected with 30% sucrose and 40 μm coronal sections cut on a cryostat. For quantitative analyses, two coronal sections per animal corresponding to -1.23 to -1.5mm AP were used. Images were acquired with an inverted fluorescence microscope (THUNDER Imaging System, Leica Microsystems) using 20x/0.8 NA objective equipped with an sCMOS camera. Image acquisition was controlled with LAS X software (Leica Microsystems). Images were acquired using 1µm optical thickness and the same imaging parameters were used for all the animals. Mosaics were generated by merging several individual frames, using a spatial overlap of 15% also performed with LAS X software. Quantification of CTB+ cells was performed manually using ImageJ on images from z stacks using DAPI and CTB staining. Analysis was performed in a blinded fashion. S1 and S2 regions of the somatosensory cortex were demarcated by the pattern of CTB back-labeling. Fifty nuclei were randomly selected using the 'multi-point tool' in layer 4 of the S1 and S2 regions. Then the images were switched to CTB channel and proportion of CTB+ cells among the 'selected' nuclei was calculated by going through images in z-stack. Data are presented as the percentage of CTB+ cells out of selected DAPI+ cells. Animals from four different litters were used.

***In vitro electrophysiology***

Acute hippocampal slices were prepared as in Basilico et al. 2019 (9). Briefly, *C3r* KO/*C3r* KO; *Thy1*::EGFP male mice at P40 were decapitated under halothane anesthesia and whole brains were rapidly immersed for 5-10 min in chilled artificial cerebrospinal fluid (ACSF: 125 mM NaCl, 2.5 mM HCl, 2 mM $CaCl_2$, 1 mM $MgCl_2$, 1.25 mM $NaH_2PO_4$, 1.1 mM glucose, 2.6 mM $NaHCO_3$) with 250 mM glycerol. Brains were sectioned into 250-µm-thick slices at 4 ˚C, using a vibratome (DSK, Dosaka EM). Slices were placed in a chamber filled with oxygenated ACSF to recover for 1 hour at room temperature (RT). All recordings were performed at RT on slices submerged and perfused with ACSF with 10 µM bicuculline. CA1 pyramidal neurons were visualized with an upright Axioscope microscope (Zeiss) and were patched in whole-cell configuration. Borosilicate glass micropipettes (3.5-4.5 MΩ) were filled with an intracellular solution (135 mM $CsMetSO_4$, 10 mM HEPES, 2 mM MgATP, 0.3 mM NaGTP, 2 mM Qx314 bromide, 2 mM $MgCl_2$, 0.4 mM $CaCl_2$, 5 mM BAPTA). Membrane currents were recorded with a patch-clamp amplifier (Axopatch 200A, Molecular Devices) and were filtered at 2 kHz, digitized (10 kHz) and acquired with Clampex 10 software (Molecular Devices). To record sEPSCs, each neuron was clamped at -70 mV for 10 minutes. Miniature EPSCs (mEPSCs) were recorded after 10 minutes of bath perfusion with tetrodotoxin (TTX, 1 μM). Recorded signals were low-pass filtered at 1 kHz and analyzed using Clampfit 10.4 software (Molecular Devices). 19-21 cells were recorded for sEPSCs and 12-14 cells were recorded for mEPSCs. sEPSC were identified on the basis of a template created for each neuron using 50-70 single events for each trace. All events recognized through the template search function were visualized, identified and accepted by manual analysis. To record evoked EPSCs, bipolar theta micropipettes (filled with ACSF) were used for stimulation and placed in stratum radiatum near CA1 area over the Schaeffer-commissural afferent fibers. Input-output curves of eEPSCs were recorded by sequentially stimulating Schaeffer collateral fibers at different intensities (0.1, 0.5, 1, 3, 7 and 10 mA) using paired-pulse protocol (0.1 ms duration of the stimulus, 50 ms interval between two consecutive stimuli and 10 s interval of two pair of stimuli). The experiments were performed from 1 to 8 hours after slicing. 19-26 cells were recorded for eEPSCs recordings. The recordings were carried out blind to animal genotype.

***Resting state fMRI***

Resting state fMRI (rs-fMRI) experiments were performed as previously described (10–12). At the time of imaging, mice were between 19 and 42 weeks old (wild-type controls: N = 19, age 30±7 weeks; *C3r* knockout: N = 20, age 28±6 weeks, animals from 13 different litters). Briefly, mice were anesthetized with isoflurane (5% induction), intubated and artificially ventilated (2% maintenance). The left femoral artery was cannulated for continuous blood pressure monitoring and terminal arterial blood sampling. After surgery, isoflurane was discontinued and replaced with halothane (0.75%), and fMRI acquisitions commenced 45 minutes after isoflurane cessation. Arterial Blood gases (pCO2 and pO2), and fluctuations of cortical BOLD-fMRI signal-to-noise ratio (SNR) were measured at the end of functional fMRI acquisitions, and compared between groups in order to discard genotype-dependent physiological confounds and anesthesia sensitivity. Mean pCO2 and O2 levels recorded in wild-type controls (22±4 and 210±32 mmHg) and *C3r* knockout (22±5 and 232±27 mmHg) showed no significant differences (2-sample t-test). Body mass and SNR also showed no significant differences between wild-type controls (28.7±5 gm and 20.8±32) and *C3r* knockout mice (28.7±4 gm and 22.1±32), excluding possible confounds from anesthesia depth in our intergroup comparisons (13).

rsfMRI images were acquired with a 7.0-T MRI scanner (Bruker Biospin, Milan) as previously described (14), using a 72-mm birdcage transmit coil and a 4-channel solenoid coil for signal reception. For each session, *in-vivo* anatomical images were acquired with a fast spin echo sequence (repetition time [TR] = 5500 ms, echo time [TE] = 60 ms, matrix 192 × 192, field of view 2 × 2 cm, 24 coronal slices, slice thickness 500 μm). Co-centered single-shot BOLD rsfMRI time series were acquired using an echo planar imaging (EPI) sequence with the following parameters: TR/TE = 1000/15 ms, flip angle 30°, matrix 100 × 100, field of view 2.3 × 2.3 cm, 18 coronal slices, slice thickness 600 μm for 1920 volumes.

***Functional Connectivity Analyses***

fMRI images were preprocessed as previously described (11, 12). The first 120 volumes were discarded to allow for T1 and gradient thermal equilibration. Data were despiked, motion-corrected, and spatially registered to an in-house common reference template. Motion traces from head realignment parameters (3 translations + 3 rotations) and the mean ventricular signal were regressed out as nuisance covariates. Denoised data was bandpass-filtered between 0.01-0.1 Hz and spatially smoothed with a full-width at half-maximum kernel of 0.6 mm.

In order to perform an unbiased investigation of brain regions exhibiting genotype-dependent connectivity alterations, we mapped Local Functional Connectivity (LFC) at the voxel-level. LFC strength here is defined as the averaged Pearson's correlation of a voxel's time-course to a subset of voxels in the local vicinity. We limited this vicinity to connections within a 6-voxel radius, and correlation coefficients were transformed to z-scores using the r-to-z Fisher's transform before averaging, and then back-transformed into correlation mean scores (13). Genotype-dependent LFC differences were assessed at the voxel-level using a 2-tailed Student's t-test and cluster corrected with the family-wise error method (FWER) as implemented by FSL ($|t| > 2$, $p < 0.05$). Quantifications of LFC at the regional level were done by averaging each subject's LFC scores within a region-of-interest (ROI) in the Prefrontal Cortex (PFC) and the mediodrsal Thalamus (TH), and inter-group differences were computed using a 2-tailed Student's t-test ($|t| > 2$, $p < 0.05$) and corrected for multiple comparisons with the Benjamini-Hochberg procedure of False Discovery Rate (FDR) $q = 0.05$. Intergroup differences in the extension and intensity of long-range rsfMRI correlation networks were mapped using a seed-based approach as previously described (15). Genotype-dependent functional connectivity differences to the prefrontal cortex were assessed at the voxel-level using a 2-tailed Student's t-test and cluster corrected with the family-wise error method (FWER) as implemented by FSL ($|t| > 2$, $p < 0.05$). Long-range connections were assessed by computing interhemispheric homotopic connectivity (correlation coefficients) between mirroring cortical Regions of Interest (ROIs, see Figure 4D), or by probing the spatial extension of fMRI connectivity of the ACC using seed-based voxelwise mapping ($|t| > 2$, $p < 0.05$) (14). Statistical significance of intergroup correlation strength in each homotopic ROI pair was assessed with a 2-tailed Student's t-test ($|t| > 2$, $p < 0.05$) and corrected for multiple comparisons with the Benjamini-Hochberg procedure of False Discovery Rate (FDR) $q = 0.05$.

##### *Statistical analysis*

Statistical analysis was performed using either Graphpad 5.0 or Sigmaplot. Plots were obtained with GraphPad Prism 5.0. Each data point refers to an individual animal. The data are presented as mean± SEM. Spine density analysis, retrograde labelling, quantification of apoptotic cells, cortical thickness and cell counts were compared using two-way ANOVA with Tukey's multiple comparison. Axon density, total axon in optic nerve, quantification of microglial engulfment of PSVue and caspase-3 labelled cells were performed with two-tailed Student's t-test or Mann-Whitney test. rs-fMRI data was analyzed by unpaired t-test. We used a 95% CI. A *p* value of $<0.05$ was set for rejecting the null hypothesis.

#### Supplementary Figures

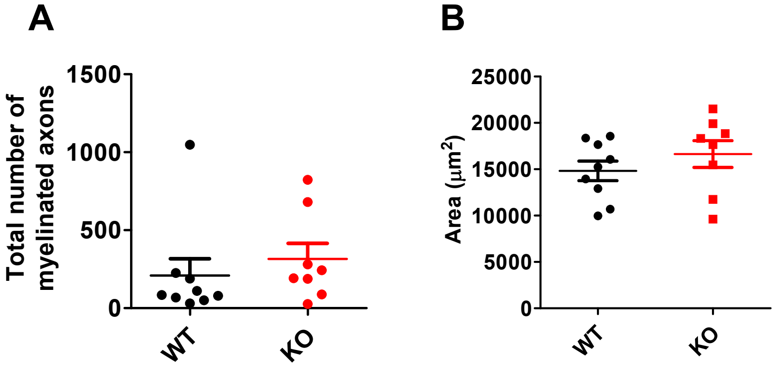


Figure S1

**Figure S1.** ***Deficient retinal ganglion cell axonal pruning in C3r knockout mice.*** (**A**) No difference in the number of myelinated axons were seen between *C3r* knockout and control littermates (unpaired t-test, p = 0.485). (**B**) A small, non-significant increase in the area of the optic nerve was observed in *C3r* knockout mice (unpaired t-test, p = 0.322; mean ± SEM, * $p < 0.05$, ** $p < 0.01$).

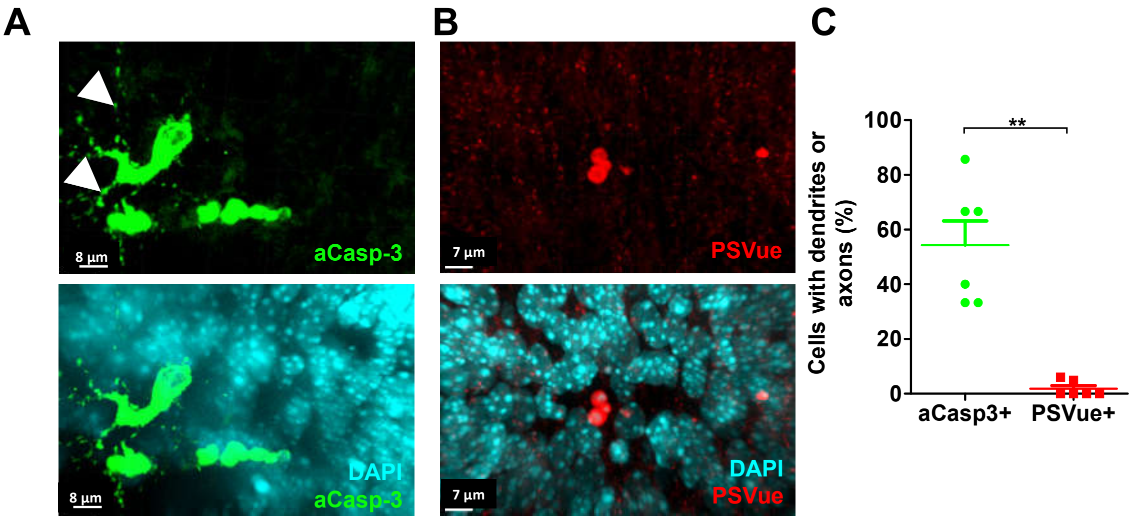


### Figure S2

**Figure S2.** ***Microglia phagocytose apoptotic neurons in early postnatal cortex.*** (**A**) Activated caspase-3 positive cells (aCasp3+) often exhibit a clear neuronal morphology with blebs of dendrites and axons. (**B**) Unlike aCasp3+ cells, PSVue+ cells do not exhibit clear neuronal morphology. (**C**) Quantitation of aCasp3+ and Phosphatidylserine exposing PSVue+ cells with dendritic branches visible (Mann-Whitney Test, $p = 0.004$; mean ± SEM, * $p < 0.05$, ** $p < 0.01$).

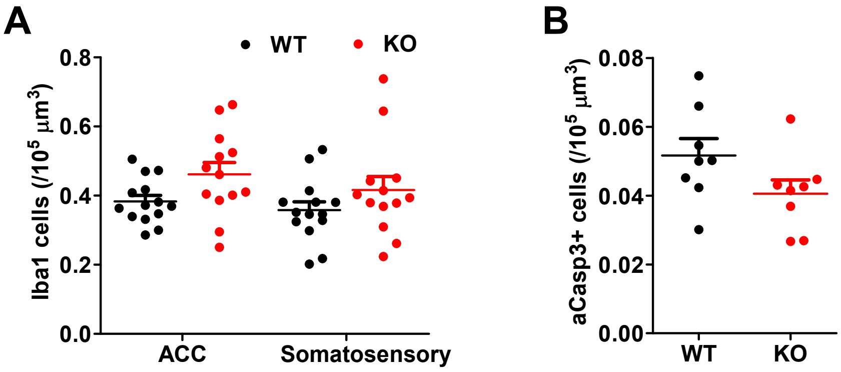


### Figure S3

**Figure S3.** ***Deficient neuronal elimination in C3r knockout mice.*** (**A**) Marginal increase in the density of microglia is seen in *C3r* knockout mice (two-way ANOVA with Tukey's post hoc test – main effect of region: $F[1, 50] = 1.46$, $p = 0.232$; main effect of genotype: $F[1, 50] = 5.44$, $p = 0.024$; region x genotype interaction: $F[1, 50] = 0.122$, $p = 0.728$). (**B**) Quantification of apoptosis initiation by aCasp3 staining revealed no differences between control and *C3r* knockout mice (unpaired t-test, $p = 0.102$)

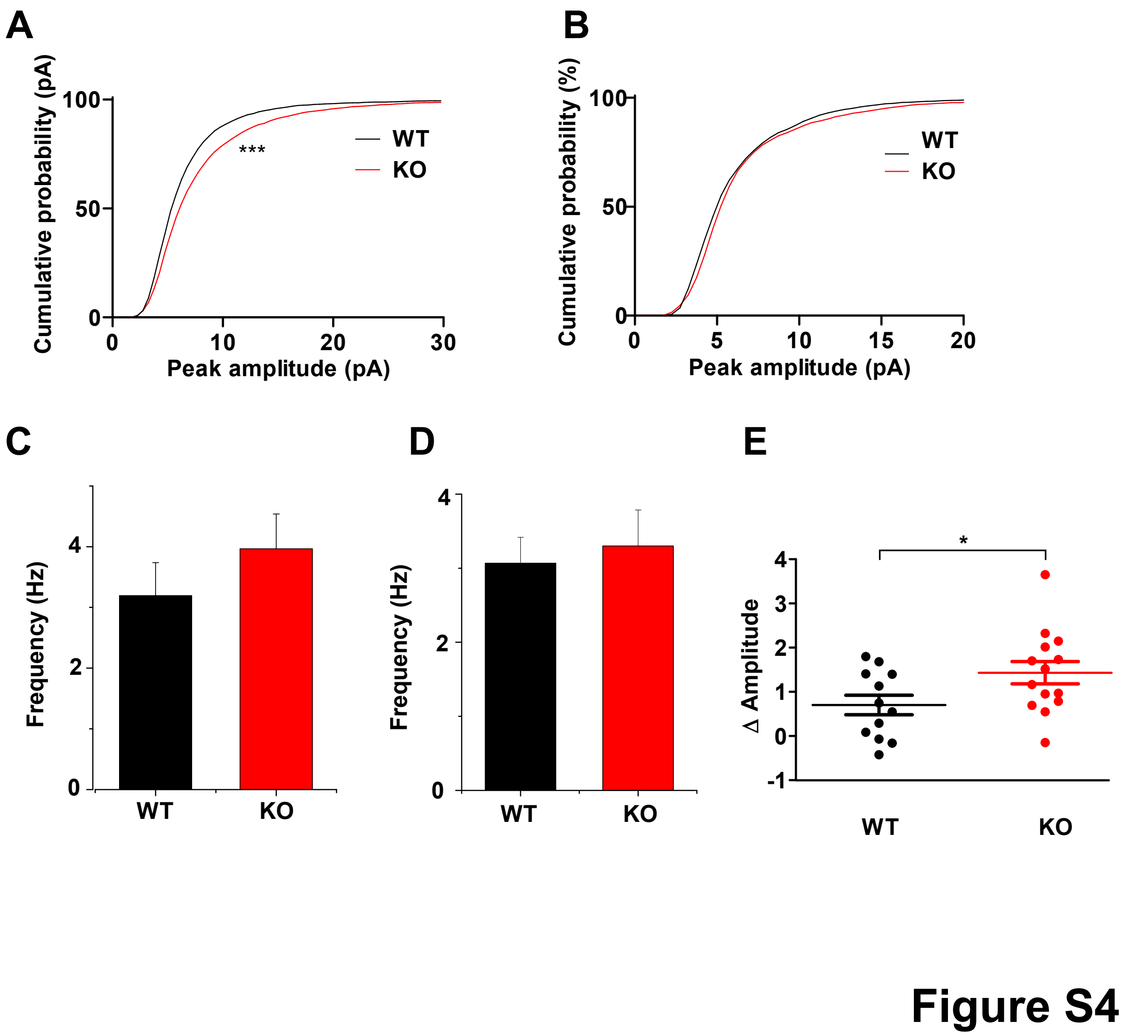


**Figure S4.** ***Increased synaptic connectivity in C3r knockout mice.*** (**A**) *C3r* knockout mice show rightward shift in amplitude of sEPSCs compared to wild -ype littermates (Kolmogorov-Smirnov test, $p < 0.001$). (**B**) No difference in the amplitude of mEPSCs were detected between the two genotypes (Kolmogorov-Smirnov test, $p = 0.0597$). (**C**) *C3r* knockout mice show no difference in sEPSC frequency compared to wild-type littermates (unpaired t-test, $p = 0.34$). (**D**) No difference in frequency of mEPSC were detected between the two genotypes (unpaired t-test, $p = 0.701$). (**E**) Mild increase in Δ amplitude (sEPSC-mEPSC) in *C3r* knockout mice (unpaired t-test $p = 0.042$; mean ± SEM. * $p < 0.05$, ** $p < 0.01$, *** $p < 0.001$).

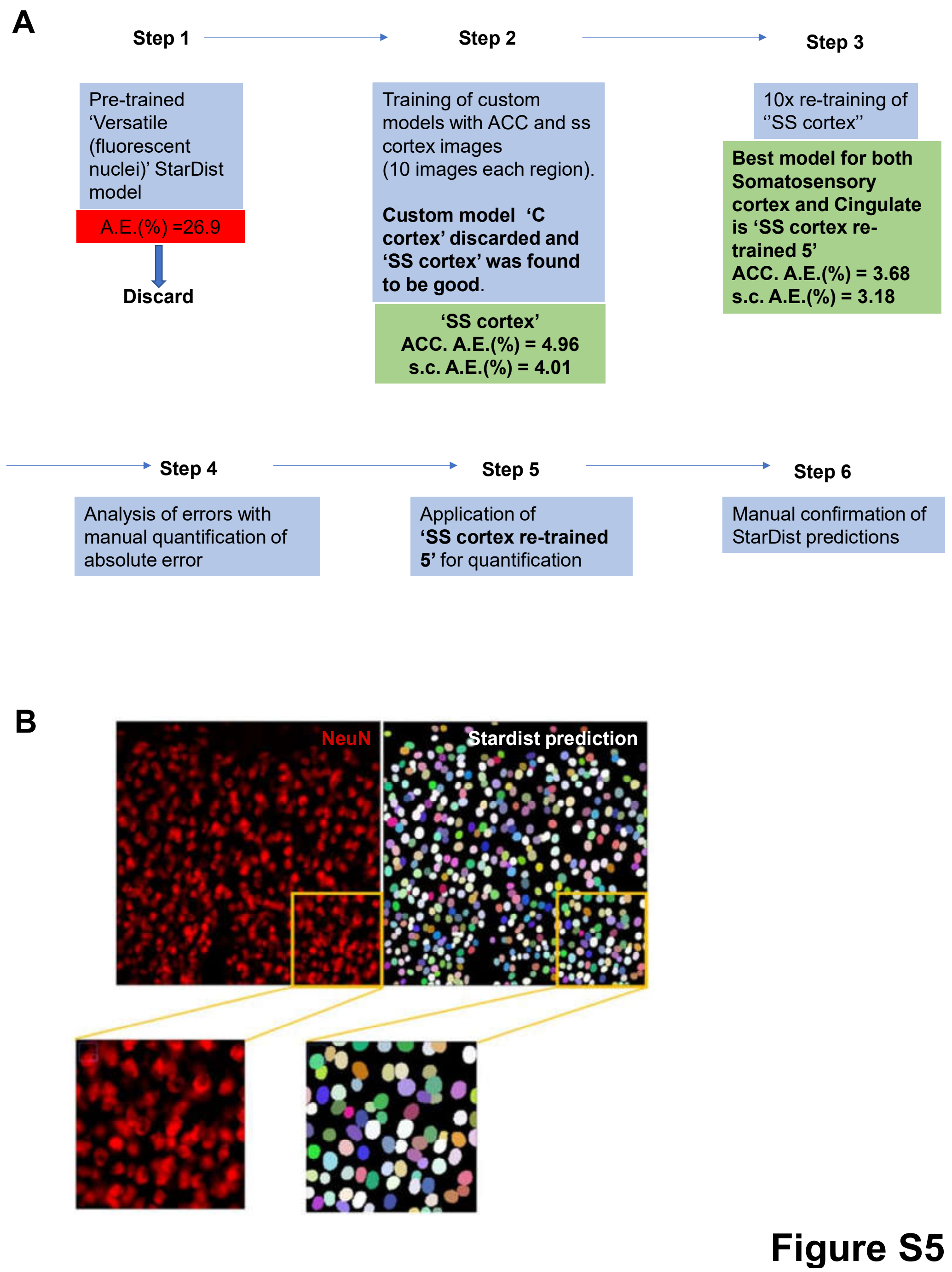


**Figure S5.** ***Development and validation of automated quantification of neurons using Stardist.*** (**A**) Flow chart depicting the development and validation of Stardist models for automated quantification of fluorescently labelled NeuN positive cells. (**B**) Example of a Stardist prediction using 'SS cortex re-trained 5' model. All the quantifications from Stardist predictions were manually verified for eliminating false positives and false negatives.
